# Human-specific features of cerebellar astrocytes and Purkinje cells: an anatomical comparison with mice and macaques

**DOI:** 10.1101/2024.08.13.607849

**Authors:** Christa Hercher, Kristin Ellerbeck, Louise Toutée, Xinyu Ye, Refilwe Mpai, Maria Antonietta Davoli, Alanna J. Watt, Keith K. Murai, Gustavo Turecki, Naguib Mechawar

## Abstract

Little is known about the morphological diversity and distribution of cerebellar astrocytes in the human brain and how or if these features differ from those of cerebellar astrocytes in species used to model human illnesses. To address this, we performed a comparative post-mortem examination of cerebellar astrocytes and Purkinje cells (PCs) in healthy humans, macaques, and mice using microscopy-based techniques. Visualizing with canonical astrocyte markers glial fibrillary acidic protein (GFAP) and aldehyde dehydrogenase-1 family member L1 (ALDH1L1), we mapped astrocytes within a complete cerebellar hemisphere. Astrocytes were observed to be differentially distributed across the cerebellar layers, displayed overall increases in area coverages with evolution, and showed features uniquely hominoid. Stereological quantifications in 3 functionally distinct cerebellar lobules demonstrated opposing trends for the canonical astrocyte markers across species with ALDH1L1+ astrocytes increasing with evolution and GFAP+ astrocytes decreasing. PC analyses revealed that while humans have the lowest PC densities, their cell body sizes were the largest with more ALDH1L1 immunoreactive astrocytes surrounding. Notably, the cognitive lobule crus I displayed the highest ratio of Bergmann glia to PC in all species. These findings align with the growing literature for astrocyte and PC heterogeneity and suggest cerebellar astrocyte and PC divergence both within and across species, possibly indicative of a role for these cells in higher-order cerebellar processing.

## Main Points

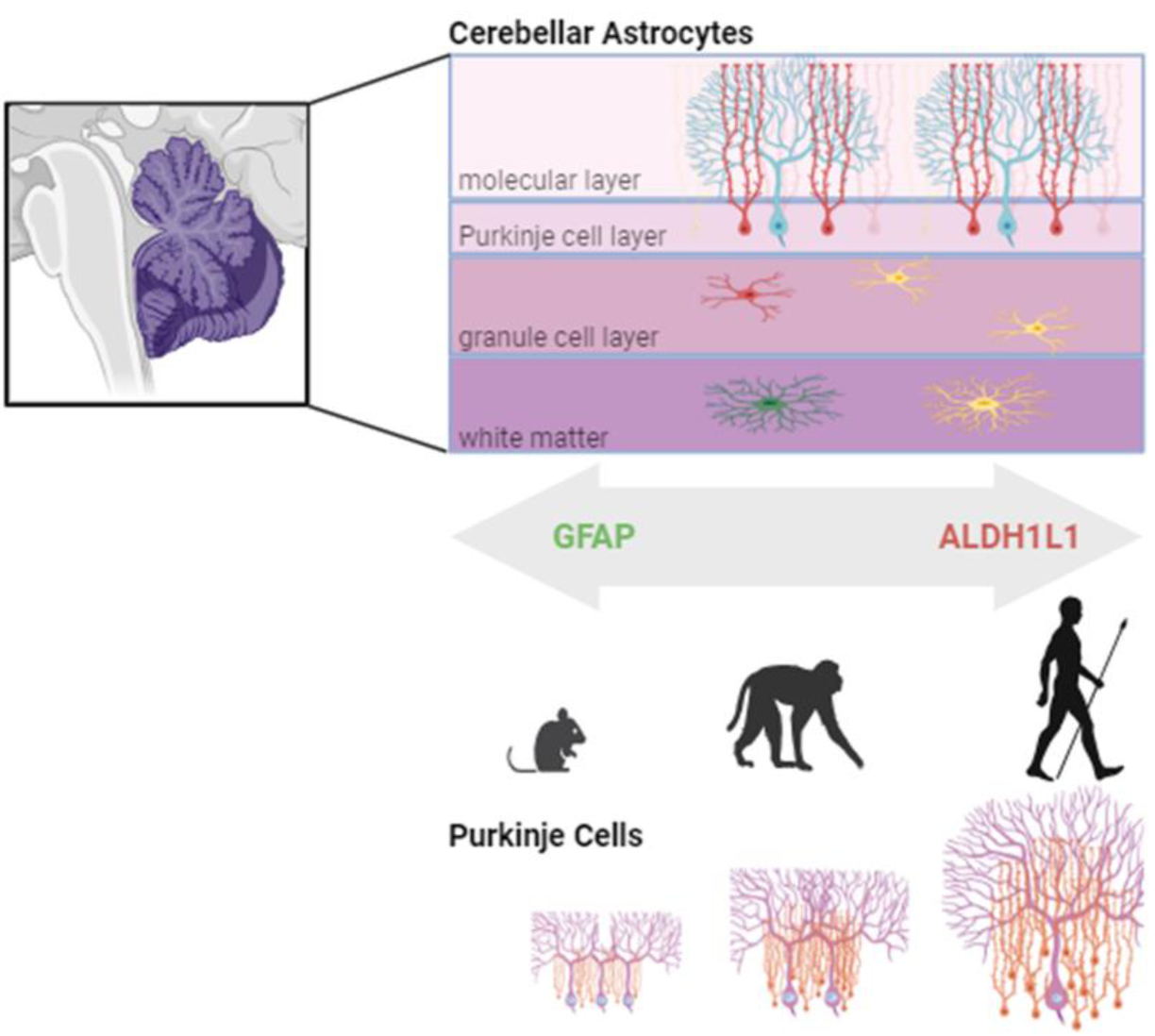

- ALDH1L1 and GFAP immunoreactive astrocytes are differentially distributed across cerebellar layers and lobules.
- Opposing trends across species with ALDH1L1+ astrocyte densities increasing and GFAP+ astrocyte densities decreasing with larger cerebella.
- With evolution, Purkinje cell densities decrease while their cell bodies and the number of Bergmann glia surrounding increase.

## 1. Introduction

More recently, it has been established that astrocytes are a highly heterogeneous glial cell population involved in a variety of roles essential for multiple biological functions such as regulating the blood brain barrier through perivascular astrocytic endfeet (Alvarez et al., 2013; Diaz-Castro et al., 2023), modulating synapses and synaptic plasticity through perisynaptic astrocyte processes (Chung et al., 2015; Stogsdill et al., 2023), and mediating the transcellular exchange of ions, second messengers, and metabolic chemicals through gap junctions (Mayorquin et al., 2018).

In the cerebral cortex, comparative neuroanatomical studies have highlighted astrocyte diversity across species, with humans exhibiting greater morphological complexity than rodents (Oberheim et al., 2006, 2009; Sosunov et al., 2014; O’Leary et al., 2020). Additionally, morphological and functional heterogeneity across species is emphasized by the presence of species-specific astrocytes such as the interlaminar astrocytes (Colombo et al., 2004; Falcone et al., 2019) and varicose projection astrocytes (Oberheim et al, 2009; Falcone et al., 2022) in the neocortex of human and non-human primates. Complementing such fine-detailed neuroanatomical reports, transcriptomic studies have further magnified the diverse signatures of astrocyte subtypes both within and between brain regions (Batiuk et al., 2020; Bayraktar et al., 2020; Endo et al., 2022; Karpf et al., 2022). However, whether species-specific features and distributions of astrocytic subtypes exist in hindbrain regions such as the cerebellum has not been well studied.

Throughout evolution, the cerebellum has expanded in parallel with the neocortex (Barton & Venditti, 2014; Sereno et al., 2020), yet its overall cytoarchitecture has remained conserved with a distinct trilaminar patterning comprising the molecular (ML), Purkinje cell (PCL), and granule cell (GCL) layers. Cerebellar astrocytes display a high degree of diversity based on the layer they reside in, which likely contributes to their functional heterogeneity (Cerrato et al., 2018; Matias et al., 2019). For example, velate astrocytes are localized in the GCL, enwrapping their processes around granule cells and cerebellar glomeruli (Hoogland & Kuhn, 2010). This positioning suggests that velate astrocytes may regulate tissue homeostasis and cerebellar circuit function (Cerrato, 2020). Spanning the PCL and ML, Bergmann glia (BG) are specialized astrocytes derived from radial glia with processes in close association with PC dendritic trees and somas. These astrocytes are involved in cerebellar development, PC synaptogenesis, and regulating synaptic activity (Buffo & Rossi, 2013). Scattered non-uniformly throughout the PCL and to varying degrees in the ML, Fañana cells are little studied and poorly understood cerebellar astrocytes (Goertzen & Veh, 2018). Finally, traditional fibrous astrocytes located in the white matter (WM) align with axons where they are thought to offer structural and metabolic support (Cerrato, 2020).

While there is growing appreciation for cerebral astrocyte heterogeneity in rodents, less is known about the morphological diversity and distribution of cerebellar astrocytes in humans. Yet this is an important question, especially given that their features may differ from those of cerebellar astrocytes in species used to model human illnesses. Thus, this study aimed to characterize cerebellar astrocytes and PCs in healthy humans and to compare them with astrocytes from non-human primate macaques and mice. Our results suggest that astrocyte and PC divergence exists, both within and across species, while also identifying species commonalities of cerebellar astrocytes across species.

## 2. Materials and Methods

### 2.1 Brain Samples

*Human cerebella:* Post-mortem human cerebella samples from 4 adults (2 males and 2 females) were obtained from the Douglas-Bell Canada Brain Bank (https://douglasbrainbank.ca) with no prior history of inflammatory, psychiatric, or neurological disorders prior to death (Table 1). *Animal cerebella:* Cerebella were acquired from 2 adult male cynomolgus macaques (generous gift from Dr. Michael Petrides) and from 2 adult male Aldh1L1-Cre/ERT2; Rosa26-TdTomato transgenic mice (generous gift from Dr. Keith K. Murai). Transgenic mice were generated by crossing Aldh1L1-Cre/ERT2 mice (JAX stock no. 031008) with Ai9 Rosa26-TdTomato reporter line mice (JAX stock no. 007909) (Madisen et al., 2010; Srinivasan et al., 2016). Breeding and animal procedures were carried out with the approval of the McGill Animal Care Committee in accordance with the Canadian Council on Animal Care guidelines.

**Table 1:** Individual Demographic Information.

| Table 1: Individual Demographic Information |  |  |  |  |
| --- | --- | --- | --- | --- |
| Human |  |  |  |  |
| Sex | Age (years) | Cause of death | PMI (hours) | pH |
| male | 54 | accidental | 54.5 | 6.3 |
| male | 36 | accidental | 49.0 | 6.3 |
| female | 54 | natural | 85.3 | 6.2 |
| female | 38 | natural | 43.0 | 5.9 |
| Cyno Macaque |  |  |  |  |
| Sex | Age (years) |  |  |  |
| male | 14 |  |  |  |
| male | 15 |  |  |  |
| ALDH1L1 Cre/ERT; Rosa26tdTomato Transgenic |  |  |  |  |
| Sex | Age (months) |  |  |  |
| male | 2 |  |  |  |
| male | 2 |  |  |  |

### 2.2 Tissue Processing

*Qualitative and % Area Coverage Assessments: Human cerebella:* 2”x 3” human sagittal cerebellar slabs comprising the complete right hemisphere were dissected at three anatomical levels, lateral X=26, deep cerebellar nuclei (DCN) X=20, and vermis X=6 (Figure 1), following the atlas of Schmahmann et al., 2000. Slabs were suspended in a 30% sucrose solution until equilibrium was reached followed by flash freezing in -35°C isopentane. For each anatomical level, cerebellar hemispheres were sectioned at 30µm in the sagittal plane, mounted on 2”x 3” glass slides, dried overnight at room temperature (RT) followed by immediate immunolabelling. *Animal cerebella*: Mice and macaques were perfused intracardially with ice-cold phosphate-buffered saline (PBS) followed by 4% formaldehyde in 0.1 M phosphate buffer. Brains were rapidly removed and postfixed in 10% neutral buffered formalin followed by separation of the cerebella from the cerebrums. Cerebella were suspended in a 30% sucrose solution until equilibrium was reached followed by rapid freezing in -35°C isopentane. Cerebellar hemispheres were systematically and exhaustively sectioned into 30µm-thick sagittal sections, mounted on Superfrost glass slides, and stored at -80°C until immunostaining.

**Figure 1.**
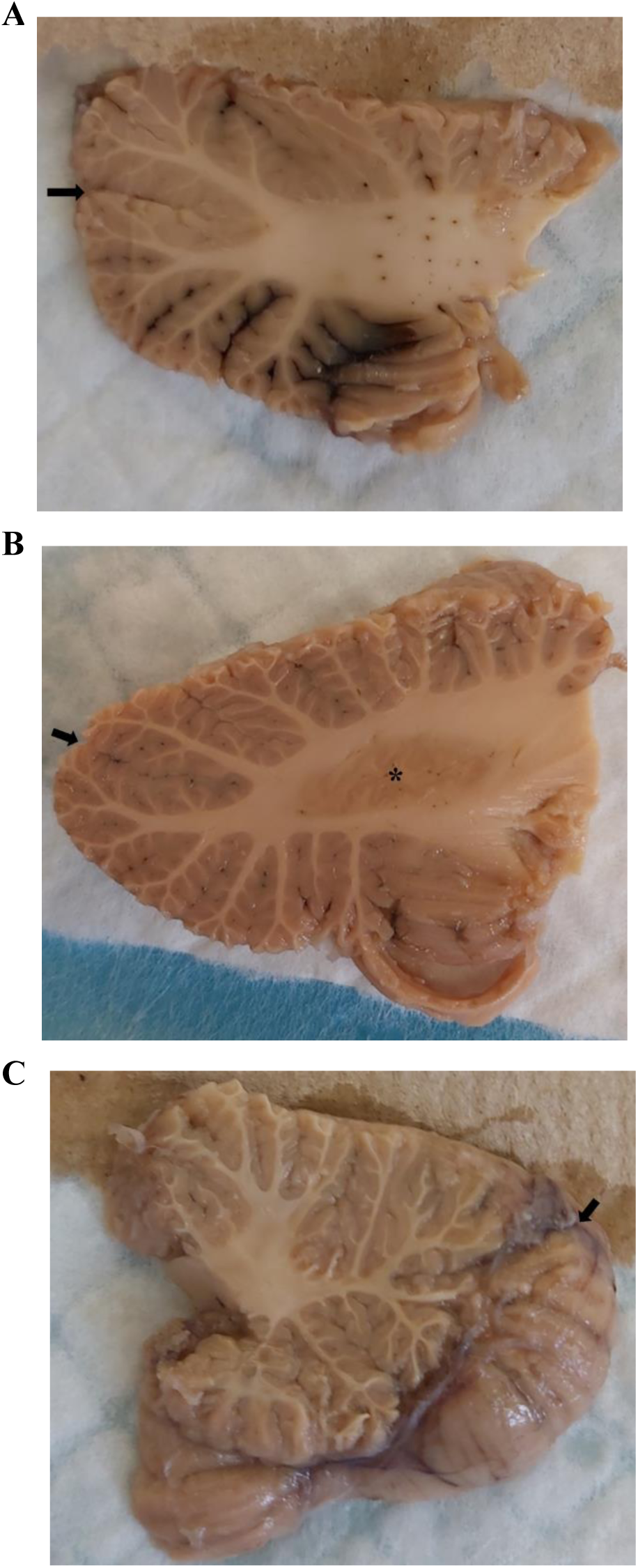
Representative sagittal slabs at 3 anatomical positions that were dissected from the human cerebellum. Right hemisphere is shown. **A** lateral, **B** deep cerebellar nuclei at the level of the dentate nucleus, **C** vermis. The lateral surface is shown in A and B. The medial surface is shown in C to display the entire vermis. Black arrows point to great horizontal fissure. Asterisk labels dentate nucleus (B).

*Stereology: Human cerebella:* 1cm^3^ tissue dissections from formalin fixed sagittal slabs were taken at the level of the dentate nuclei (lobule III and crus I, X=20, Schmahmann et al., 2000) and in the vermis (lobule VIIA folium, X=6, Schmahmann et al., 2000) and prepared as above. These lobules were chosen as each represents a unique functional domain: the vermis associating with emotional regulation, crus I with cognition, and lobule III with traditional cerebellar motor functions (Buckner et al., 2011; Schmahmann, 2019). 1cm^3^ human tissue blocks were then systematically and exhaustively sectioned into 30µm-thick sagittal sections, mounted on Superfrost glass slides, and stored at -80°C until immunostaining. An equally spaced series of sections per lobule of interest were immunolabelled (4-7 sections/subject/anatomical position).

*Animal cerebella:* Three series of sections were sampled based on anatomical position, one containing lobule III, one encompassing crus I, and a final series was taken throughout the vermis, guided by corresponding atlases (Allen Mouse Brain Atlas; Madigan & Carpenter, 1971). In mice and macaques, 3-7 sections/animal/region of interest were immunolabelled. Section sampling for all species followed systematic and random sampling principles of stereology (Mouton, 2002; Kreutz & Barger, 2018).

### 2.3 Immunolabeling

Sections were air-dried for 7 minutes and heated in a 60°C oven for either 15 minutes (mice and macaques) or 30 minutes (humans) to aid in section adherence to the glass slides. Sections were cooled for 5 minutes at RT followed by rinsing in PBS for 5 minutes. Antigen retrieval was performed using proteinase K (1:1000) for 15minutes followed by PBS washes. Sections were blocked in 10% normal donkey serum, (NDS, Jackson ImmunoResearch, 017-000-121) and PBS + 0.2% Triton-X (mice and macaques) or 10% NDS and PBS (humans) for 1 hour at RT. Mouse sections were incubated overnight at 4°C in 10% NDS + PBS + 0.2% Triton-X with chicken anti-GFAP (1:250, abcam ab4674). Macaque sections were incubated for 48 hours at 4°C in 10% NDS + PBS + 0.2% Triton-X with chicken anti-GFAP (1:250, abcam ab4674) and mouse anti-ALDH1L1 (1:50 millipore MABN495). Human sections were incubated overnight at 4°C in 10% NDS + PBS with chicken anti-GFAP (1:1000, abcam ab4674) and mouse anti-ALDH1L1 (1:250, Millipore MABN495). Following primary antibody incubations, sections were rinsed in PBS followed by application of secondary antibodies, Alexa Fluor® 488-conjugated donkey anti-chicken (1:500, Jackson ImmunoResearch, 703-545-155) (applied to all species) and Alexa Fluor® 647-conjugated donkey anti-mouse (1:500, Jackson ImmunoResearch, 715-605-151, macaque and human sections) for 1 hour at RT. Following PBS washes, sections were quenched for autofluorescence using TrueBlack® (Biotium, 23007) for 75 seconds, rinsed and coverslipped with Vectashield Vibrance with DAPI mounting medium (Vector, H-1800).

### 2.4 Image Analysis

*Qualitative and % Area Coverage Assessments:* Three sections/species were sampled representative at each anatomical position: lateral, DCN level, and vermis level. An additional 3 sections/species, adjacent to the immunofluorescent sections, were stained for cresyl violet (Nissl stain) acting as a histological guide. Slides were scanned on an Evident Scientific VS120 slide scanner at 20X. Exposure times remained constant for all sections within one species. Cerebellar layers, ML, PCL, GCL, and WM, in all cerebellar lobules were outlined and annotated using QuPath (Bankhead et al., 2017, version 0.3.2) guided closely by reference atlases for mice (Allen Mouse Brain Atlas), macaques (Madigan & Carpenter, 1971), and humans (Schmahmann et al., 2000). For the human cerebellar sections, 1 complete folia per lobule was chosen for annotations in order to maximize efficiency (Figure 2). Percent area coverages for GFAP-IR and ALDH1L1-IR astrocytes in each layer of each lobule were obtained using a threshold approach based on the mean intensity and standard deviation of our signal of interest. Pixels at and above the threshold value were considered positive while pixels below the threshold were reflected as background (Figure 3).

**Figure 2.**
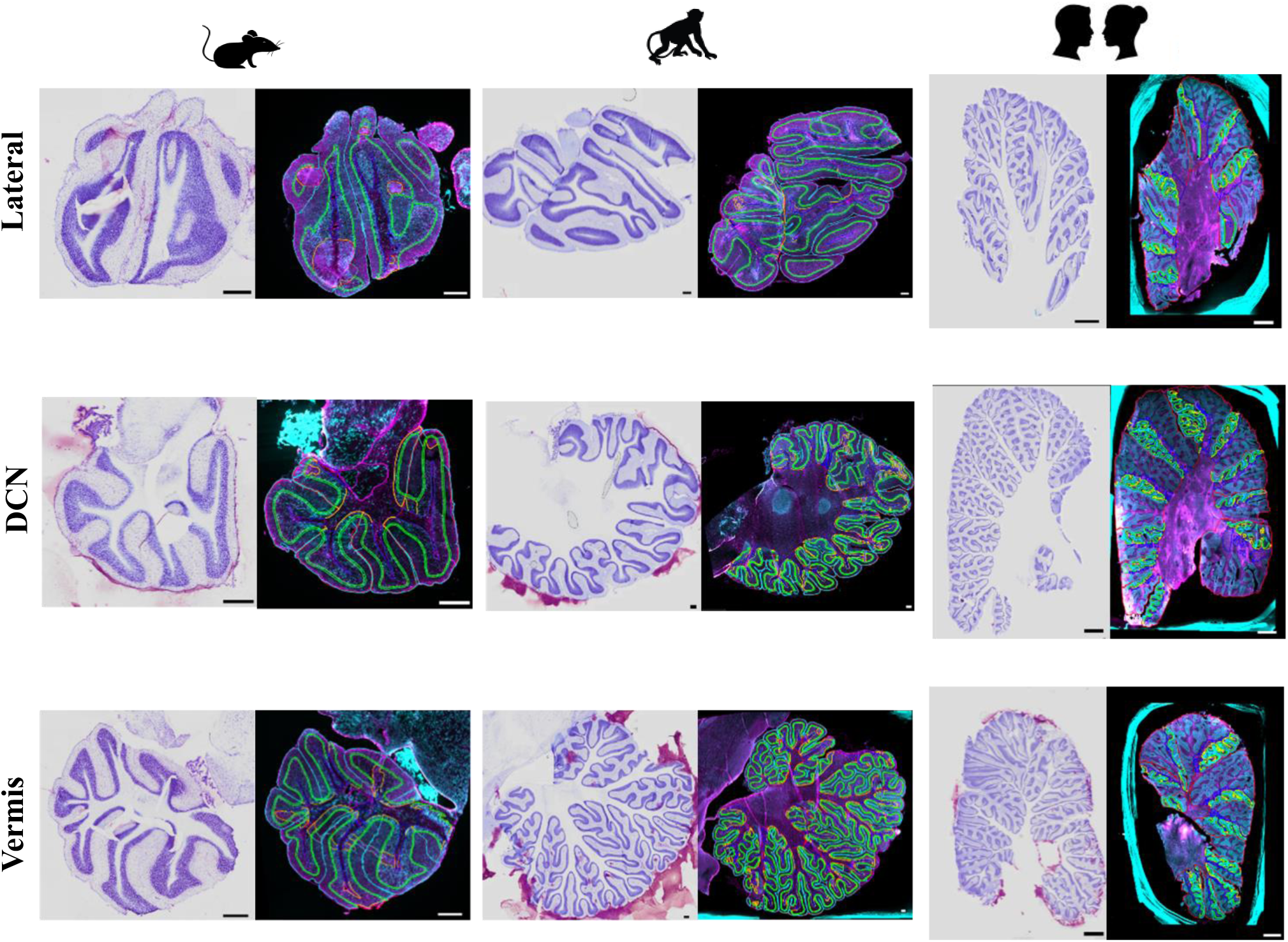
Representative images at 3 anatomical positions (lateral, DCN, vermis) that were analyzed in the mouse, macaque, and human cerebellum. Left image displays nissl stained section for histological reference. Right image displays immunolabelling for GFAP astrocytes in magenta and ALDH1L1 astrocytes in cyan. Lobules and layers were annotated and outlined using QuPath. DCN deep cerebellar nuclei. Mouse and macaque scale bar 500µm, human scale bar 5mm.

**Figure 3.**
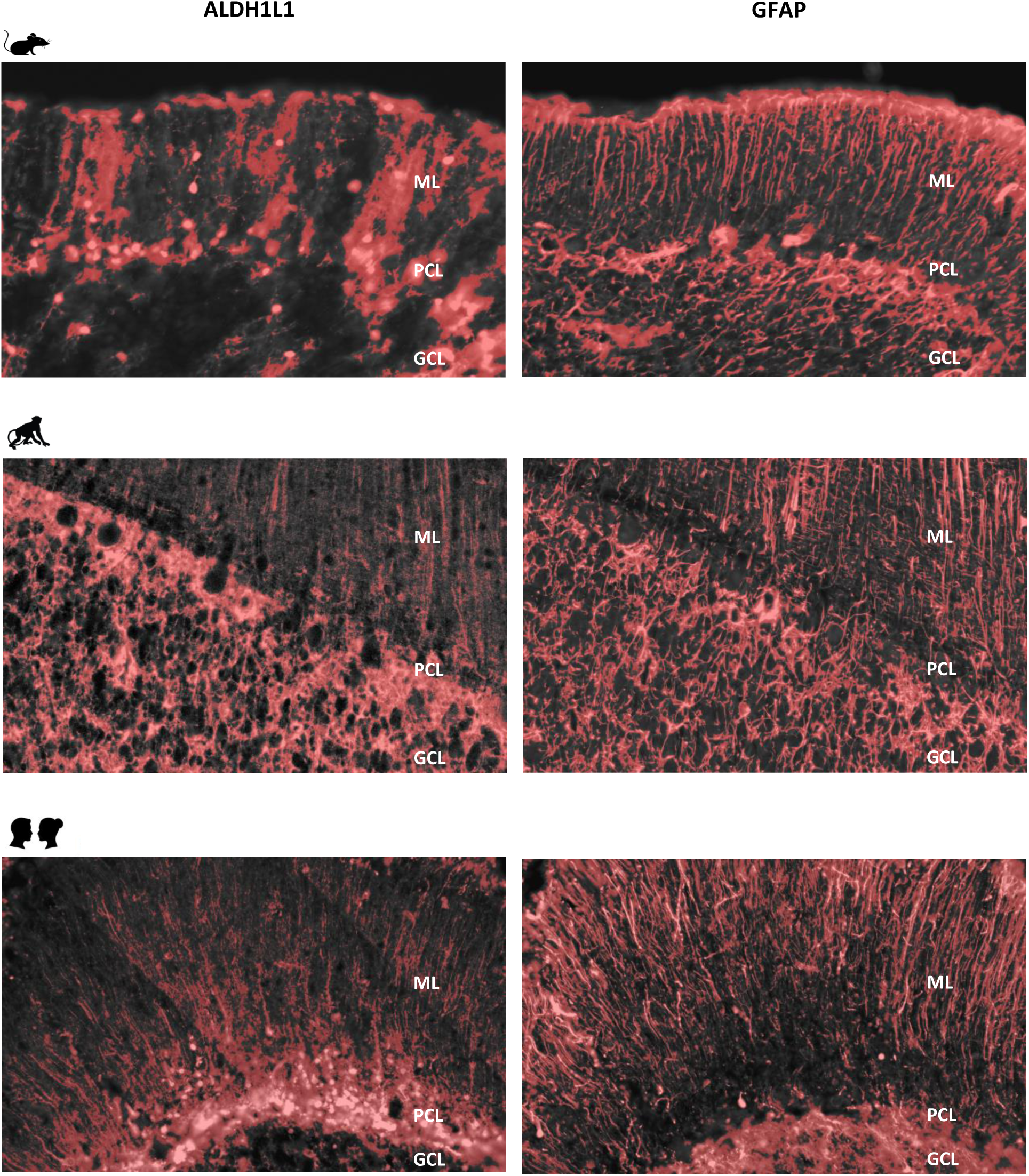
Threshold representations of % area coverages in the mouse, macaque, and human cerebellum for ALDH1L1 and GFAP immunoreactive astrocytes. Red overlay indicates pixels included in the analysis which were based on identifying the mean intensity and standard deviation of our signals of interest. ML molecular layer, PCL Purkinje cell layer, GCL granule cell layer.

*Stereology Assessments:* Unbiased stereology was performed using the software StereoInvestigator (MBF bioscience, United States). Image stacks were acquired every 2μm throughout our mounted tissue thickness using a Zeiss ApoTome2 Axio Imager.M2 microscope system at 63X (N.A 1.4) in each lobule and layer of interest for individual species. Tissue thickness was measured at every sampling site to account for wavy tissue and to ensure optimal image quality. The optical fractionator probe was applied to the image stacks using a 9μm dissector height with 1μm guard zones. Unbiased estimates of ALDH1L1^+^ and GFAP^+^ astrocytes were quantified in the PCL, GCL, and WM. In the PCL, the nucleator probe (isotropic sampling, 4 rays) was applied to the image stacks where for each individual species, 60 PC body and nucleus sizes were measured in each lobule of interest. PCs were counted directly using the optical fractionator probe in the PCL to minimize imaging times. Volume estimates were generated by applying the Cavalieri probe on the same contours used for counting in each layer. The robustness of our stereological estimates was indicated by obtaining coefficients of error (Gunderson m=1) <0.10 (Supplemental Table 1).

## 3. Results

### 3.1 Differential immunoreactivity of canonical astrocyte markers GFAP and ALDH1L1 across cerebellar layers

To understand the similarities of cerebellar astrocytes in human tissue compared to model animals including mice and primates, we obtained tissue from all 3 species (Figure 1) and performed quantitative analysis of astrocytes at three different positions in the cerebellum: lateral, at the level of the DCN, and in the vermis (Figure 2). We used both ALDH1L1 and GFAP as markers for astrocytes (Figure 3), since each marker labels different subsets of astrocytes. We found in all three species that the ML of the cerebellum was largely comprised of GFAP+ processes with few astrocytic cell bodies. The complexity of these processes increased with species evolution, with humans displaying more branched and horizontal processes compared to mice and macaques (compare Figures 4a, 5a, 6a). Varicosity-like protrusions and knotted blebbing’s along these processes were also observed in some of the human samples (Figure 6a inset). In mice and macaques, GFAP+ bulbous endfeet were strikingly visible near the upper limit of the ML (Figure 4a, 5a). The PCL contained mainly ALDH1L1+ cell bodies consistently observed across species (Figure 4b, 5b, 6b), with these astrocytes being in close proximity to PCs. Despite the density of granule cells, velate astrocytes were highly visible across species in the GCL (Figure 4c, 5c, 6c). In mice and humans, ALDH1L1+ astrocytes presented with a dense cell body labelling (Figure 4c & 6c) whereas macaques displayed a punctate labelling (Figure 5c). Characteristic fibrous astrocytes were observed in the WM with GFAP-IR being particularly intense in these cells across species (Figure 4d, 5d, 6d). Aligning with a previous report in the cerebral cortex (O’Leary et al., 2020), we also noted astrocytic twin cells in human cerebellar WM (Figure 6d inset). Within the DCN, we observed robust species differences in astrocytes. There was a dramatic absence of GFAP-IR astrocytes in the DCN of mice (Figure 4e) which was observed across all nuclei (interposed, dentate, and fastigial). In contrast, ALDH1L1 was observed in high abundance across these nuclei in mice (Figure 4e), confirming the presence of astrocytes in mouse DCN. In macaque DCN, astrocyte processes were highly visible and appeared discretely organized (Figure 5e). Astrocyte cell bodies in both macaque and human DCN were difficult to distinguish from processes (Figure 5e, 6e).

**Figure 4.**
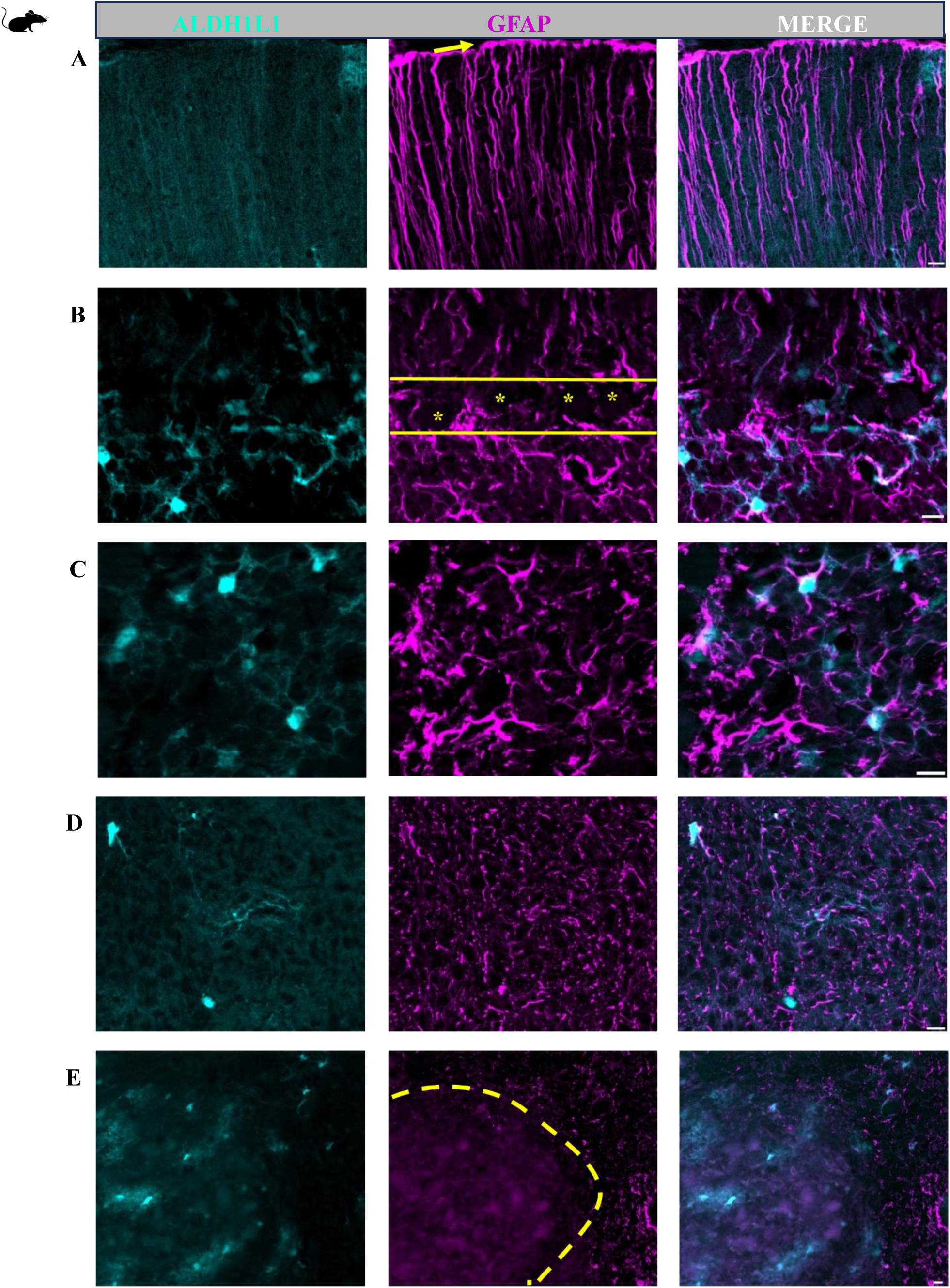
Representative images of cerebellar astrocytes in the mouse. **A** molecular layer, **B** Purkinje cell layer, **C** granule cell layer, **D** white matter, **E** deep cerebellar nuclei (fastigial nucleus is shown). Bulbous astrocytic end feet were observed at the upper limit of the molecular layer (A yellow arrow). Purkinje cell layer is outlined by solid yellow with Purkinje cells (B yellow asterisk’s). In the granule cell layer astrocyte cell bodies showed robust ALDH1L1+ labelling while GFAP was highly visible in astrocytic processes (C). Note the scarcity of GFAP in the deep cerebellar nuclei (E). Yellow dashed line outlines deep cerebellar nuclei (E). Scale bar 10µm.

**Figure 5.**
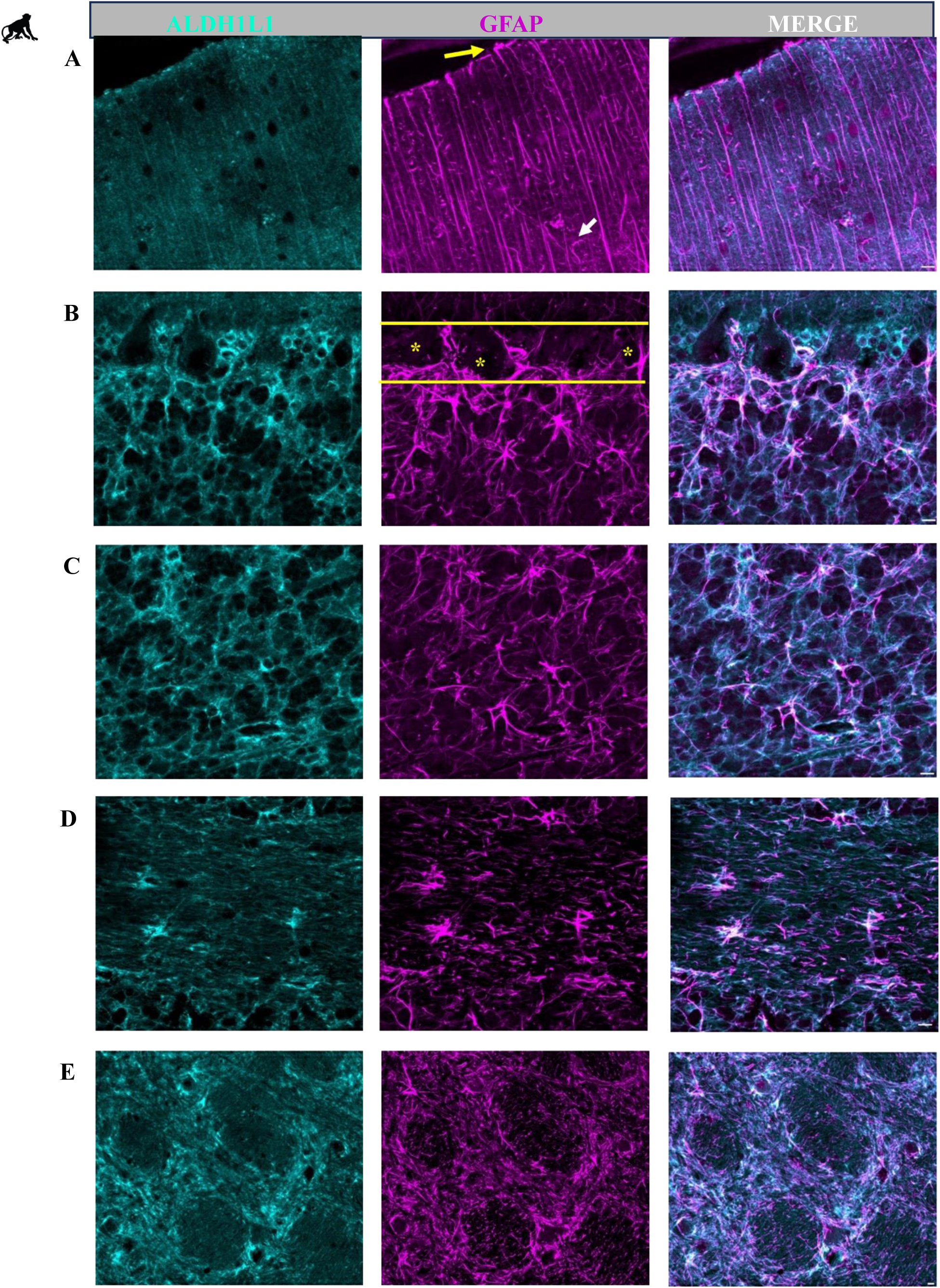
Representative images of cerebellar astrocytes in the macaque. **A** molecular layer, **B** Purkinje cell layer, **C** granule cell layer, **D** white matter, **E** deep cerebellar nuclei (fastigial nucleus is shown). Similar to mice, bulbous astrocytic end feet were observed at the upper limit of the molecular layer (A yellow arrow). The presence of lateral appendages on Bergmann glia processes were also observed (A white arrow). Yellow solid line outlines the Purkinje cell layer and yellow asterisk’s labels Purkinje cells (B). Astrocytes in the granule cell layer displayed a mesh-like patterning (C). Typical fibrous astrocytes were observed in the white matter (D). In the deep cerebellar nuclei ALDH1L1+ and GFAP+ astrocyte cell bodies and processes showed a discrete organization (E). Scale bar 10µm.

**Figure 6.**
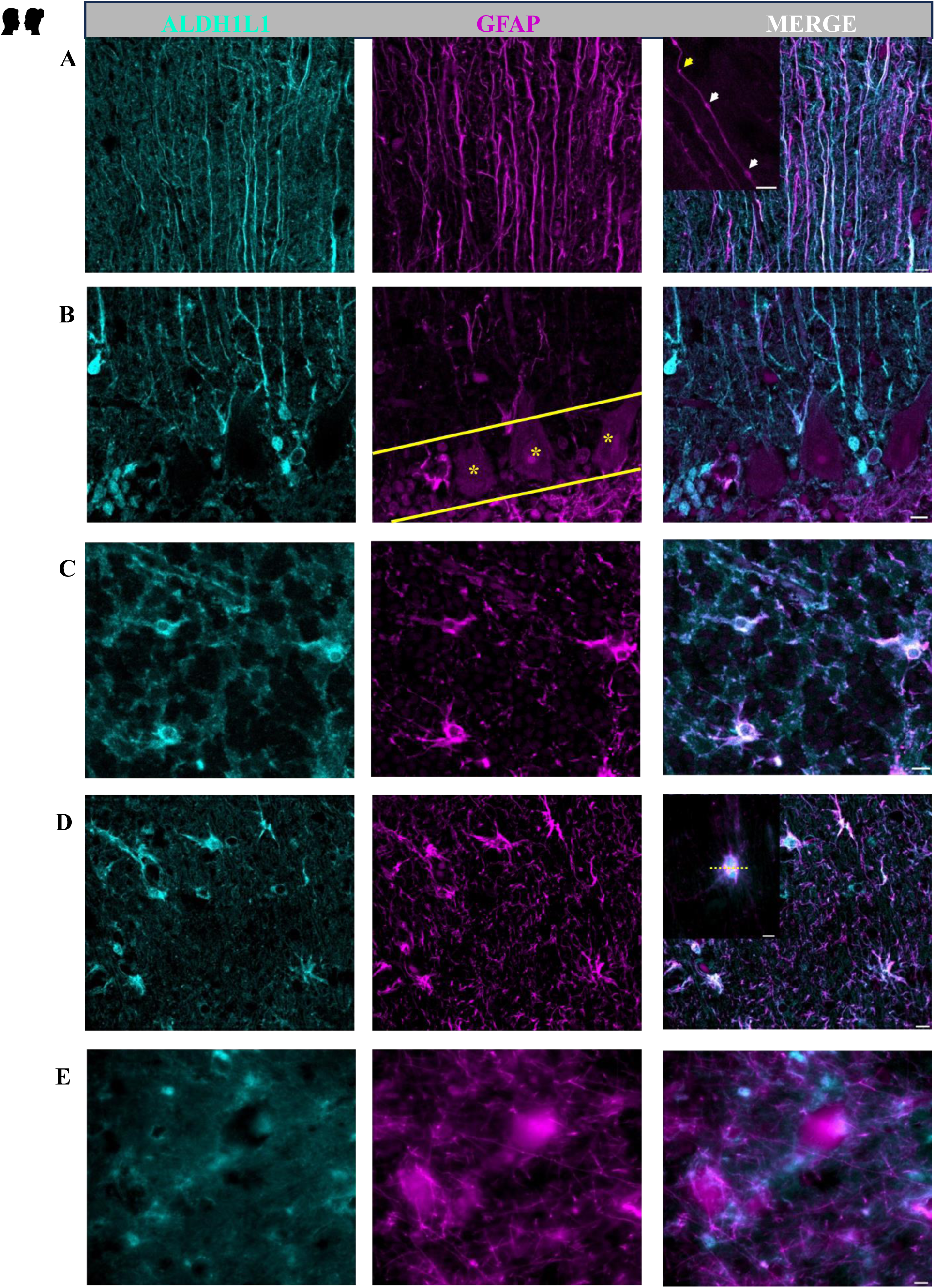
Representative images of cerebellar astrocytes in humans. **A** molecular layer, **B** Purkinje cell layer, **C** granule cell layer, **D** white matter, **E** deep cerebellar nuclei (fastigial nucleus is shown). In the molecular layer, ALDH1L1+ and GFAP+ processes displayed complex horizontal branching with lateral appendages highly visible (A). Along GFAP+ processes, knotted-blebbings were visible (inset A white arrows) as well as varicosity like protrusions (inset A yellow arrow). In the Purkinje cell layer, ALDH1L1 strongly labelled Bergmann glia (B). Yellow solid line outlines Purkinje cell layer and yellow asterisk’s labels Purkinje cells (B). Velate astrocytes were observed around granule cells and presumed cerebellar glomeruli in the granule cell layer (C). Fibrous astrocytes were highly visible in the white matter (D). Twinning of white matter astrocytes was also observed (inset D yellow dotted line divides two astrocytes). In the deep cerebellar nuclei astrocyte cell bodies were difficult to distinguish (E).

Overall, these qualitative observations indicate layer-specific preferences for astrocytic markers. Furthermore, species-specific differences were apparent with respect to ALDH1L1 and GFAP immunolabeling, particularly in the PCL and DCN respectively.

### 3.2 Astrocytic % area coverages increase with species evolution

We next wondered whether the area fraction of astrocytes increased with species complexity. To address this, we obtained % area coverages for ALDH1L1+ and GFAP+ cerebellar astrocytes (Figure 7). Comparing the cerebellum in its entirety, we observed that the % area coverage of GFAP+ astrocytes was increased slightly in macaques and humans (33%) compared to mice (30%). However, the % area coverage of ALDH1L1+ astrocytes increased steeply with species evolution, with 22% in mice, 24% in macaques, and 35% coverage in humans. It is well established that functionally distinct astrocyte subtypes reside in specific cerebellar layers with BG positioned in the PCL, velate astrocytes in the GCL, and fibrous astrocytes in the WM. Therefore, we were interested in exploring % area coverages of our astrocyte markers in these layers. We observed variations in immunoreactivity, with species-specific patterns. In the ML, GFAP+ astrocyte % area coverages were slightly increased in macaques (33%) and human (33%) compared to mice (29%), similar to overall cerebellar values. Surprisingly, ALDH1L1 % area coverages were similar between mice (23%) and humans (24%) and lowest in macaques (14%). The similarity between mice and humans could be due to the use of transgenic mice with TdTomato labelling astrocytes in their entirety in comparison to antibodies which may miss detecting astrocyte processes where the protein labeled is low, thus likely underreports the area occupied by astrocytes in macaques and humans. We observed the greatest species differences in the PCL with the % area coverage of ALDH1L1+ astrocytes being 35% in mice, 42% in macaques, and 54% in humans. Interestingly, while ALDH1L1 coverage was highest in the PCL, the % area coverage of GFAP+ astrocytes in the human cerebella was slightly reduced in human cerebellar PCL (24%) compared to mice and macaques (29%). In the GCL, GFAP+ astrocyte % area coverages were slightly increased in humans and macaques (34%) compared to mice (29%). Similar values were observed for the % area coverages of ALDH1L1+ astrocytes where we observed increases in humans (32%) compared to mice and macaques (24%). In WM, we observed robust species differences for both astrocyte markers. The % area coverage of GFAP+ astrocytes was 31% in mice, 38% in macaques, and 43% in humans. The % area coverage of ALDH1L1+ astrocytes in mice was strikingly low (13%) compared to macaques (21%) and humans (29%). We found that there were striking species differences in the DCN, in which the coverage of ALDH1L1+ astrocytes was 19% in mice compared to 29% in both macaques and humans. The coverage of GFAP+ astrocytes in the DCN was profoundly reduced in mice with only 6% in mice compared to 38% and 42% coverage in macaques and humans, respectively. These observations were similar across the dentate, interposed and fastigial DCN (Figure 7).

**Figure 7.**
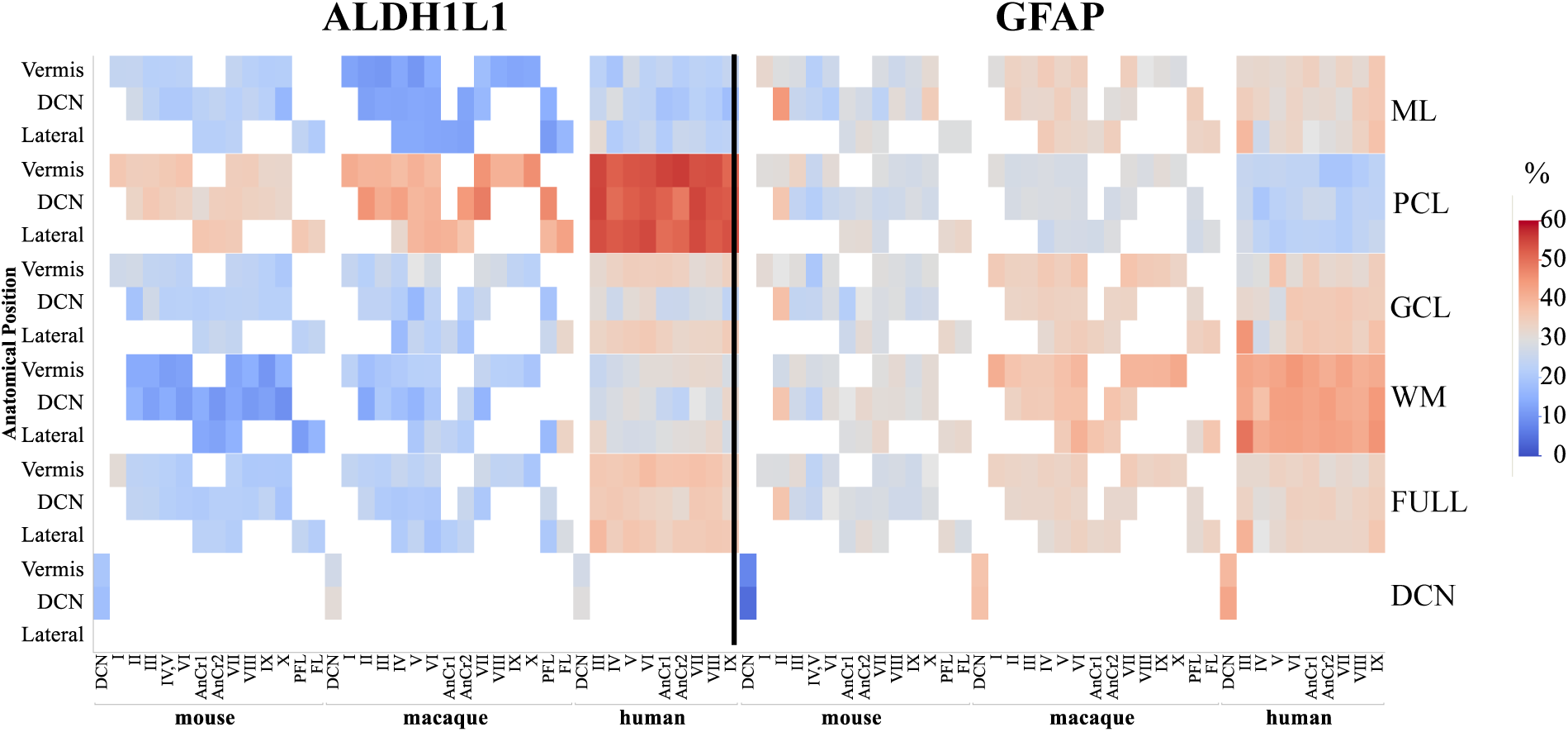
Area coverage of ALDH1L1+ and GFAP+ astrocytes in cerebellar layers and lobules in mice, macaques, and humans (percentages are reported). Note the high expression of GFAP+ astrocytes in the molecular layer and in the white matter in comparison to ALDH1L1+ astrocytes. ALDH1L1+ astrocytes were highly expressed in the Purkinje cell layer, especially in the human cerebellum. GFAP+ astrocytes in the deep cerebellar nuclei in mice displayed very low coverage compared to macaques and humans. ML molecular layer, PCL Purkinje cell layer, GCL granule cell layer, WM white matter, DCN deep cerebellar nuclei. Full indicates average of all cerebellar layers excluding the DCN.

Overall, we observed differences in GFAP and ALDH1L1 astrocyte coverages across most anatomical regions of interest (lateral, DCN, and vermis) within each species (Supplemental Table 2). ALDH1L1 displayed the highest % area coverage in the PCL for all species, suggesting a high expression of this astrocytic marker in BG, whereas the lowest coverages were in WM for mice and ML for macaques and humans. GFAP % area coverages were highest in the WM for all species indicating that GFAP is a robust marker for cerebellar fibrous astrocytes. The lowest coverages were in the GCL for mice and macaques and in the PCL for humans. The largest species differences were observed in the PCL for ALDH1L1 % area coverages and in the DCN for GFAP coverages.

### 3.3 Astrocyte area coverages are partially conserved across species in cerebellar lobules

We next evaluated ALDH1L1 and GFAP % area coverages in all cerebellar lobules across species. In the mouse, lobule II (DCN level) presented the highest GFAP coverages (37%) while lobule IV/V (vermis level) displayed the lowest (24%). ALDH1L1 coverage was highest in lobule I (31%, vermis level) and lowest in lobule X (20%, DCN level). In macaques, lobule VII had the highest GFAP coverages (36%, vermis level) while lobule IV had the lowest (31%, lateral level). The flocculus displayed the highest ALDH1L1 coverages (29%, lateral level) with lobules VI and X showing the lowest (20%, lateral and vermis levels, respectively). In humans, lobule III displayed the highest GFAP coverages (41%, lateral level) while lobule IV showed the lowest (30%, lateral level). Similarly, ALDH1L1 coverages were highest in lobule III (39%, lateral level) and lowest in AnCrII (32%, DCN).

Overall, within each species, we observed subtle differences in astrocytic coverages across the cerebellar lobules. Across species, while there was divergence observed in astrocytic coverages in specific cerebellar lobules, we also observed convergence; for example, lobule IV displayed the lowest GFAP % area coverages in mice, macaques, and humans (Figure 7, Supplemental Table 2).

Current knowledge suggests lobule specific functions within the cerebellum (Buckner, 2013; Klein et al., 2016) where lobules I-V are classified as anterior exhibiting traditional motor functions while posterior lobules (VI-X) are predominantly involved in non-motor functioning. We explored whether the current astrocyte markers differed according to these functional domains and observed subtle layer and species-specific differences in GFAP and ALDH1L1 % area coverages between functional domains (anterior vs posterior) and anatomical positions (lateral, DCN, vermis) of the cerebellum (Supplemental Table 3).

### 3.4 Opposing trends for ALDH1L1 and GFAP astrocyte densities in the cerebellar cortex with species evolution

Building on the above knowledge that ALDH1L1 and GFAP % area coverages display lobule-, layer-, and species-specific heterogeneity, we next aimed to quantify these cells utilizing robust stereological principles. Based on our qualitative observations, we chose to focus our quantifications in cerebellar layers as it was difficult to visualize astrocyte cell bodies in DCN. Overall, cerebellar estimates revealed that ALDH1L1+ astrocyte densities were approximately x higher in humans compared to macaques and mice, whereas GFAP+ astrocyte densities were 1.1x lower than macaques and 2.8x lower compared to mice (Figure 8a). Within specific cerebellar layers, ALDH1L1+ astrocyte densities in the PCL were approximately 1.2x higher in humans compared to macaques and mice. An opposite pattern was observed for GFAP+ astrocyte densities where humans had 1.6x lower densities than macaques and 5x lower densities than mice. In the GCL, ALDH1L1+ astrocyte densities were approximately 2x higher in humans compared to macaques and 1.4x higher than mice. GFAP+ astrocyte densities in humans were 1.2 x higher than macaques yet 1.9x lower than mice. In WM, ALDH1L1+ astrocyte densities in humans were similar to those in macaques and mice while GFAP+ astrocyte densities were 1.3x lower than macaques and 2.2x lower than mice. Proportion of ALDH1L1+ and GFAP+ astrocytes in cerebellar layers across species. Aligning with our qualitative and % area coverage data, we found that most astrocytes in the PCL were ALDH1L1+ whereas astrocytes in the WM were largely GFAP+ (Figure 8b).

**Figure 8.**
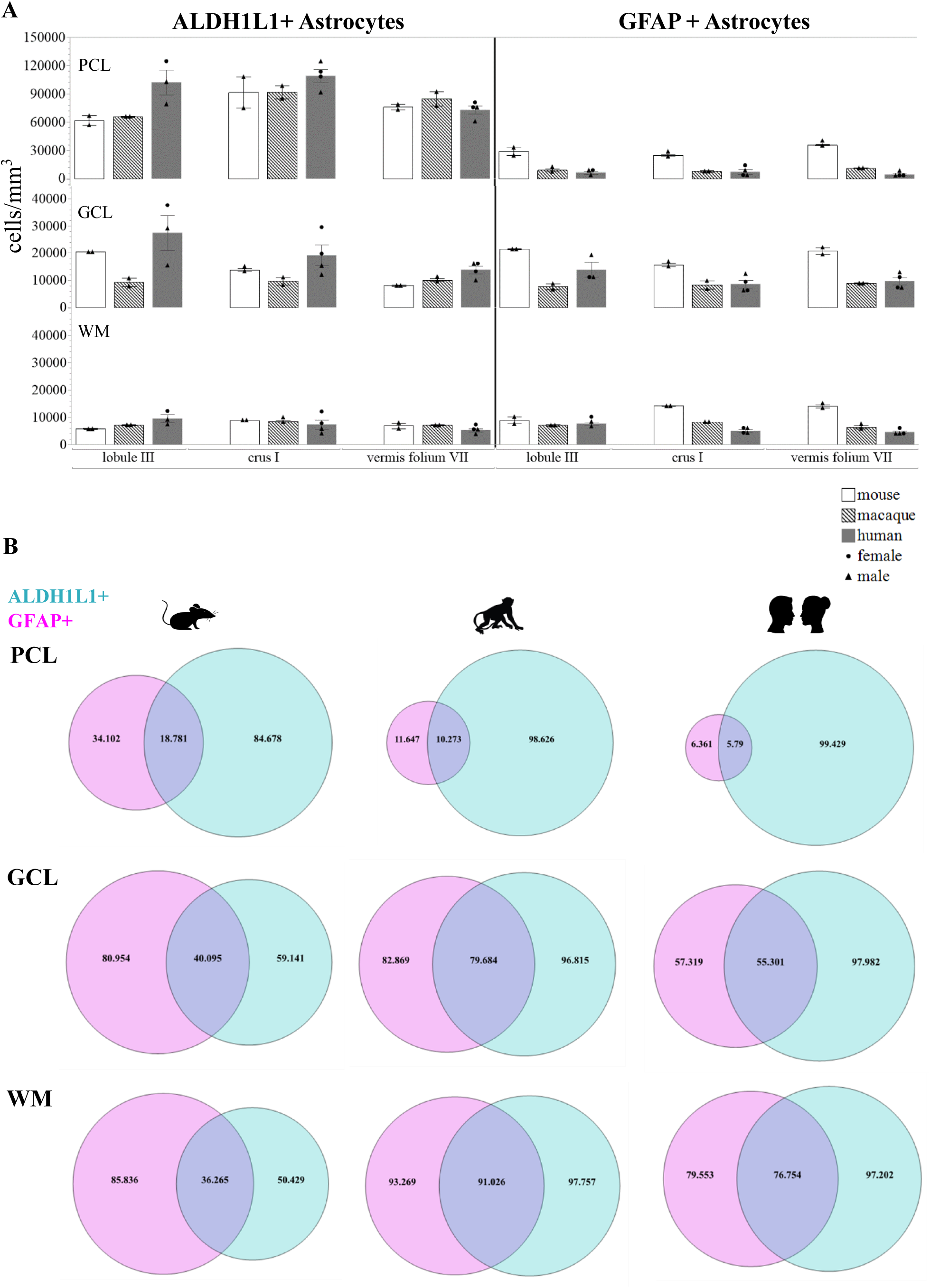
A) ALDH1L1+ and GFAP+ astrocyte densities in cerebellar layers and functionally distinct lobules of interest in mice, macaques, and humans. In all species, ALDH1L1+ astrocytes had the highest densities in the PCL and the lowest in the WM. When comparing distinct lobules in the human cerebellum, both astrocytic markers showed the lowest densities in the vermis. Opposing trends were observed across species between the astrocytic markers with ALDH1L1+ astrocytes increasing with species complexity while GFAP+ astrocytes decreased from mice to macaques to humans. **B)** Proportion of ALDH1L1+ and GFAP+ astrocytes in cerebellar layers across species. For all species, most astrocytes in the PCL were ALDH1L1+ whereas astrocytes in the WM were largely GFAP+. Most astrocytes in the human cerebellum were ALDH1L1+ whereas the majority of astrocytes in the mouse cerebellum were GFAP+. ML molecular layer, PCL Purkinje cell layer, GCL granule cell layer, WM white matter.

We quantified ALDH1L1+ and GFAP+ astrocytes in 3 functionally distinct lobules of interest (lobule III, crus I, and vermis VIIA folium) in the PCL, GCL, and WM across species (Figure 8a). In the PCL, the largest species differences were observed in lobule III where ALDH1L1+ densities in humans were 1.6-1.7x higher than macaques and mice respectively. In contrast, the vermis displayed the largest species differences for GFAP+ densities where humans had 2.5x lower densities compared to macaques and 8x lower densities than mice. In the GCL, the largest species differences were observed in lobule III where ALDH1L1+ densities in humans were 3x those of macaques and in the vermis where ALDH1L1+ densities in humans were 1.7x higher than mice. GFAP+ densities were 1.8x’s higher in humans compared to macaques in lobule III while the vermis displayed the largest difference between humans and mice with the former having 2.2x lower densities. In WM, ALDH1L1+ astrocyte densities differed the most in the vermis between humans and macaques where humans had 1.4x lower densities. ALDH1L1+ densities in lobule III showed the largest differences between humans and mice where humans had 1.7x higher densities than mice. Crus I showed the largest differences in GFAP+ densities between humans and macaques where humans had 1.7x lower densities.

Similar to the PCL and GCL, the vermis presented the largest difference between humans and mice with the former having 3.1x lower GFAP+ densities.

Overall, the human vermis displayed the lowest densities of ALDH1L1+ and GFAP+ astrocytes across all layers analyzed, which was not observed in other species. We also observed opposing trends with ALDH1L1+ densities increasing with species evolution, particularly in the PCL and GCL while GFAP+ densities decreased from mice to macaques to humans (Figure 8a). We found that most astrocytes in the human cerebellum were ALDH1L1+ whereas the majority of astrocytes in the mouse cerebellum were GFAP+ (Figure 8b).

### 3.5 GFAP-defined territories are larger in the human cerebellum

Recent reports have suggested that regions displaying smaller astrocyte territories compensate with higher astrocyte densities (Viana et al., 2023). Therefore, it was of interest to explore whether density and size were related in our datasets, particularly for GFAP+ astrocytes. Combining our measurements of % area coverage together with stereological estimates for astrocyte densities, we calculated the % area coverage per GFAP+ astrocyte yielding 2D estimates of GFAP+ defined territories (Figure 9). GFAP was chosen as the astrocytic marker of choice as it provides extensive labelling of the astrocyte processes compared to ALDH1L1. In the ML, presumed to contain largely BG processes, GFAP+ territories in humans were 3.2x larger than macaques and 10.4x larger when compared to mice. In the GCL, humans displayed territories 3.1x larger than mice. Human and macaque territories were similar in this layer. In WM, GFAP+ territories in humans were 1.8x greater than macaques and 3.7x greater compared to mice.

**Figure 9.**
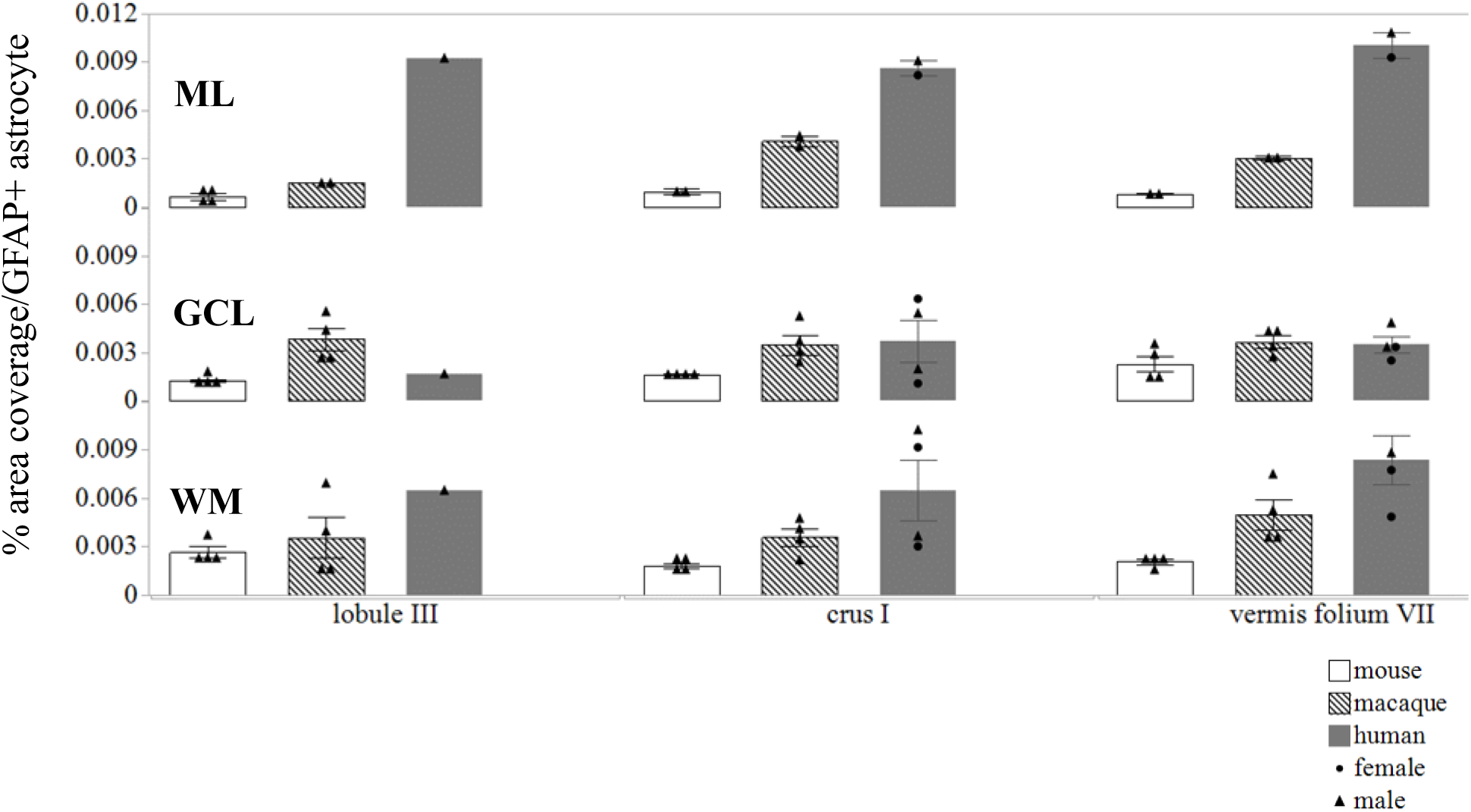
GFAP-defined astrocytic territories in cerebellar layers and functionally distinct lobules of interest in mice, macaques, and humans. Human cerebellar astrocyte territories were 3 times larger than macaques and 10 times larger than mice in the ML highlighting the complexity of Bergmann glia processes in humans. ML molecular layer, GCL granule cell layer, WM white matter.

Aligning with our previous analyses, we were interested in comparing GFAP territories in functionally distinct lobules across species (Figure 9). In the ML, the largest difference between humans and macaques was observed in lobule III with humans displaying 6.2x larger territories. When comparing human GFAP territories to mice, the vermis displayed the greatest differences with human GFAP territories being 12.4x greater than mice. A slightly different pattern emerged in the GCL where humans showed 2.1x lower territories than macaques in lobule III. However, crus I displayed the greatest species difference between humans and mice with humans having 3.8x larger territories. Similarly in WM, crus I showed the greatest species differences with human territories being 2.2x greater than macaques and 4.7x larger than in mice.

Overall, while we observed lower GFAP+ astrocyte densities in humans compared to mice and macaques, the territories of these astrocytes were larger in the human cerebellum, consistent with previous descriptions of cerebral astrocytes (Oberheim et al.,2006, 2009; O’Leary et al., 2020) (Figure 9).

### 3.6 Purkinje cell parameters diverge in mice, macaques, and humans

With the knowledge that PCs are the main inhibitory output neurons of the cerebellum that are tightly connected with BG (De Zeeuw & Hoogland, 2015), we next sought to quantify PCs in lobule III, crus I, and in the vermis lobule VIIA folium in mice, macaques, and humans (Figure 10). Overall, irrespective of lobule, PC densities in humans were 1.7x lower compared to macaques and 8.1x lower than mice (Figure 10a). However, cell body sizes were 1.2x larger in humans compared to macaque and 4.2x when comparing to mice (Figure 10b). Humans displayed an average of 14 BG surrounding 1 PC compared to 7BG to 1PC in macaques and 2 BG to 1PC in mice (Figure 10c). Focusing on the 3 functionally distinct lobules, PC densities displayed the greatest species differences in the vermis lobule VIIA folium where humans had 2.2x lower densities than macaques and 10.8x lower densities than mice (Figure 10a). With respect to cell body size, crus I showed the greatest differences between humans and macaques where humans had 1.5x larger cell body sizes (Figure 10b). Lobule III displayed the greatest differences between humans and mice with humans having 4.7x larger cell body sizes (Figure 10b). Across all species, crus I showed the highest BG to PC ratios compared to lobules III and vermis lobule VIIA folium with humans displaying an average of 17 BG surrounding 1 PC compared to 8BG to 1PC in macaques and 2 BG to 1PC in mice (Figure 10c).

**Figure 10.**
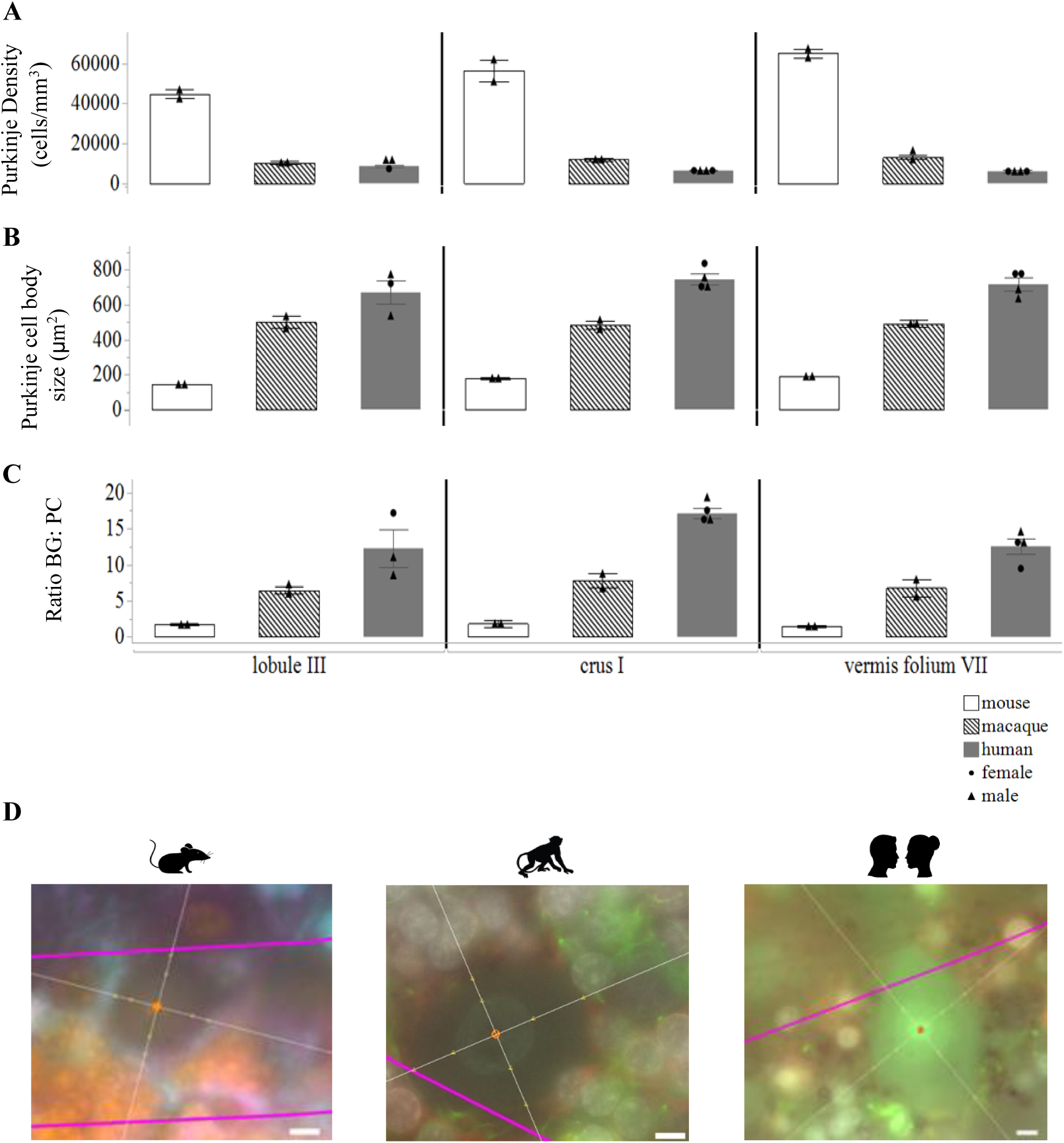
Purkinje cell parameters in mice, macaques, and humans. **A)** Purkinje cell density. Purkinje cell densities in humans were almost 2x lower than macaques and 8x lower than mice. **B)** Purkinje cell body area. Purkinje cell body sizes were 1.2x larger in humans compared to macaques and 4x bigger than mice. **C)** Ratio of Bergmann glia to one Purkinje cell. Humans had an average of 14 BG surrounding 1 PC compared to 7BG to 1PC in macaques and 2 BG to 1PC in mice. The highest ratio was in crus I, particularly in the human cerebellum. **D)** Representative examples of the nucleator probe emitting 4 rays to measure Purkinje cell body size. BG Bergmann glia, PC Purkinje cell. Scale bar 5µm.

Overall, while PC densities decreased with species evolution, cell body sizes increased as did the number of BG surrounding each PC with the greatest ratio of BG to PC being in crus I.

## 4. Discussion

In this study we performed a comparative neuroanatomical investigation to thoroughly characterized cerebellar astrocytes and PCs in healthy mice, macaques, and humans. We first mapped astrocytes within a complete cerebellar hemisphere at 3 anatomical positions and observed differential immunoreactivity of canonical astrocyte markers, GFAP and ALDH1L1 across the cerebellar layers. We then measured % area coverages of our astrocyte markers reporting increases in astrocyte coverages with evolution. Next, we quantified astrocytes in 3 functionally distinct cerebellar lobules and identified opposing trends in astrocyte markers across species with ALDH1L1+ astrocytes increasing with evolution and GFAP+ astrocytes decreasing. Finally, we performed an in-depth analysis of PCs which showed that as species complexity increased, PC densities decreased while cell body sizes increased, with increasing numbers of BG surrounding individual PCs. Notably, the highest BG:PC ratio was observed in the cerebellar lobule associated with cognitive processing.

Our qualitative observations suggest that the morphology of human cerebellar astrocytes is more complex compared to less evolved species, in line with observations made previously for neocortical astrocytes (Oberheim et al., 2009; O’Leary et al., 2020). GFAP-IR astrocytes displayed thick bulbous end-feet in the molecular layer, prominently in mice and macaques.

While the function of these end-feet has yet to be fully characterized (De Zeeuw & Hoogland, 2015), it is tempting to speculate that they are involved in the homeostasis of the blood brain barrier. Varicosity-like protrusions and knotted blebbings on BG processes were also observed in humans. Further analyses are needed to determine if these features are present in all individuals or if they are solely observed under specific conditions as has been previously reported (Falcone et al., 2022). A striking observation was the scarcity of GFAP-IR astrocytes in the DCN of mice, which has been previously reported (Shibuki et al., 1996). This observation appeared specific to the marker GFAP as ALDH1L1, glutamine synthetase, and sox9 were all visualized in the DCN (unpublished data) suggesting a scarcity of intermediate filaments in DCN astrocytes in healthy mice. An absence of intermediate filaments in astrocytes was reported to have no major consequences on nonreactive astrocytes, perhaps due to other cytoskeleton components, such as actin filaments compensating to support the development and maintenance of astrocytic processes (Wilhelmsson et al., 2004).

Complementing these qualitative observations were our robust stereological estimates for ALDH1L1+ and GFAP+ astrocytes in 3 functionally distinct lobules. In agreement with our % area coverages, ALDH1L1 was highly expressed in the PCL which was consistent across species. Interestingly, we observed opposing trends between our two astrocyte markers, with ALDH1L1+ densities increasing with species evolution, particularly in the PCL and GCL while GFAP+ densities decreased from mice to macaques to humans. This opposing trend could imply that astrocyte cellular properties may differ across species. For example, it suggests a higher metabolic demand in human astrocytes as ALDH1L1 is a key enzyme in folate metabolism which is important in nucleotide biosynthesis and cell division (Krupenko, 2009; Yang et al., 2011). Overall, human astrocytes displayed 2-5x lower GFAP+ densities than mice depending on the layer of interest, which was somewhat unexpected. GFAP is an intermediate filament protein involved with structural stabilization of astrocyte processes (Yang & Wang, 2015), and thus might be expected to increase with the complexity of processes. However, Viana et al., (2023) recently reported in mice, that while the hippocampal layers, CA1 radiatum and DG molecular, showed the lowest astrocyte densities, these layers compensated by displaying the largest defined territories. Our data directly aligns with this report where we observed lower GFAP+ astrocyte densities in the human cerebella, compared to mice and macaques, yet the territories of human astrocytes were 3-10x larger depending on the layer the astrocytes resided in. This is consistent with cerebral astrocytes displaying increased complexity in the human brain (Oberheim et al.,2006, 2009; O’Leary et al., 2020). Previous reports (Oberheim et al., 2009; O’Leary et al., 2020; Verkhratsky et al., 2021) have observed astrocytes, particularly protoplasmic astrocytes, to be arranged in discrete orientations, with non-overlapping territories tiling across the cerebral cortex, however this was difficult to assess in the current study based on our astrocyte labelling. We can speculate that like neocortical astrocytes, tiling in the WM is limited as we observed twinning of astrocytes in this layer implying process overlap. In the PCL, BG processes radiate throughout the ML often displaying lateral appendages in human and non-human primates as observed in the current study and by others (Muñoz et a., 2021). Furthermore, BG processes are highly dynamic within the ML, interacting with both excitatory and inhibitory synapses (De Zeeuw & Hoogland, 2015) making it unclear if discrete territories are maintained. There remains the possibility for independent astrocytic territories in the GCL with velate astrocytes not only encompassing granule cells but also veiling their processes around individual cerebellar glomeruli. However, future investigations are needed to visualize cerebellar astrocytes in their entirety to address if tiling is present and if there are species differences in tiling overlap, for example.

Finally, we also analyzed PCs as these output neurons are highly interconnected with BG (Yamada et al., 2000; Miyazaki et al., 2017). Previous reports have characterized PCs across species and, in accordance with the present study, PC density decreases accompanied by significant increases in cell size and spacing of PCs have been described with increasing evolution (Lange 1975; Korbo & Andersen, 1995). A recent study elegantly compared PCs in humans and mice, finding that human PCs display higher dendritic complexities and present 7.5x more dendritic spines along dendrites, processing 6.5x more input patterns than in mice PCs, suggesting greater computational capabilities in humans (Masoli et al., 2024). To our knowledge, our present study is the first to describe a BG to PC relationship in the human cerebellum. Interestingly, we observed an increase with evolution in the number of BG surrounding one PC with the highest ratio being in crus I compared to lobule III and vermis lobule VIIA folium. Considering crus I is involved in cognitive processing, our results may thus be indicative of an increased need for BG in higher level cognitive cerebellar functioning, suggesting greater computational processing in the human cerebellum (Masoli et al., 2024).

Few studies have interrogated lobule and layer specificities of PC and astrocytic distributions. Kozareva et al., (2021) used transcriptional profiling to characterize PC diversity across 16 lobules in the mouse cerebellum. They found that PCs displayed considerable heterogeneity, particularly in posterior lobules, uvula (lobule IX) and nodulus (lobule X). Although we did not perform PC analyses in these lobules, it is interesting to note that crus I (also a posterior lobule) displayed considerably larger cell body sizes and higher BG to PC ratios compared to lobule III (anterior lobule) and to the vermis lobule in humans. The authors also report regional specialization of BG in lobule VI, the uvula (lobule IX) and nodulus (lobule X). Such findings together with our observations highlight the importance of performing detailed lobule and layer assessments to parcellate the cellular heterogeneity observed within the cerebellum.

This study is not without limitations. First, caution needs to be taken when comparing ALDH1L1-IR astrocytes in mice, as the transgenic Aldh1L1-Cre/ERT2; Rosa26-TdTomato mice displayed a more complete staining pattern with intense labeling of cell bodies and astrocyte processes that likely skews the % area coverages when compared to antibody labeling. Intuitively, labeling human and macaque astrocytes in their entirety would reveal even greater species differences and thus our results likely reflect conservative estimates in these species. Also, while we attempted to define GFAP territories, the 2D method we used was rudimentary as we did not perform an in-depth Sholl analysis which would have allowed for 3D representations. However, labeling astrocytes in their entirety in the post-mortem human brain is challenging.

While vimentin is a promising marker for visualizing the complexities of astrocytes, we observed limited labeling in the cerebellum (1-2 vimentin+ astrocytes/section in humans, unpublished data). Such observations are in line with those reported in other subcortical structures such as the thalamus, where minimal vimentin+ astrocytes have been observed (O’Leary et al., 2020). Finally, we recognize that our sample sizes were limited. While we included male and female human individuals and preliminarily observed no obvious differences, increasing our sample would allow us to confidently disentangle potential sex differences in PC and astrocytes within the cerebellum.

Despite these limitations, we present the first comprehensive examination of cerebellar astrocytes and PCs in mice, macaques, and humans, highlighting features unique to humans. This original study extends the growing literature of astrocyte heterogeneity to the cerebellum and highlights species divergence in PC and astrocytic features important for our understanding of animal models of disease and translational of findings from these species to human patients. Furthermore, this study is timely considering the increasing evidence for the cerebellum in higher order cognitive and emotional brain functioning (Buckner 2013; Fastenrath al., 2022) where growing literature demonstrates aberrant structure and function of the cognitive-affective cerebellum in neurodevelopmental (Haldipur et al., 2022; Hwang et al., 2022) and psychiatric disorders (Depping et al.,2018; Romer et al., 2018; Chambers et al., 2022). Thus, this study provides foundational knowledge for cerebellar astrocyte and PC properties unique to the healthy brain, allowing future directions to next determine dysfunction of these properties in central nervous system disease and other disorders.

## AUTHOR CONTRIBUTIONS

CH and NM conceptualized and designed the study. GT participated in the acquisition and clinical characterization of the human brain samples. CH, KE, LT, RM, XY and MAD performed histological experiments. CH, KE, LT and XY conducted data analysis. AW and KM provided significant conceptual input and discussion of results. CH and NM prepared the manuscript and all authors critically revised and approved its final version. **Corresponding Author**: Naguib Mechawar, PhD, Douglas Mental Health University Institute, 6875 Blvd LaSalle, Verdun, Qc, H4H 1R2, Canada

## ACKNOWLEDGMENTS

We wish to express our heartfelt gratitude to the families of the donors who graciously agreed to donate the brains of their beloved family members. We also wish to thank Dominique Mirault, Vanessa Larivière, and Elizabeth Desouza for their skillful technical assistance with brain dissections. We thank Veronika Zlatkina and Sarah Lefebvre from Dr. Michael Petrides’ laboratory, who provided macaque tissues. The present study used the services of the Molecular and Cellular Microscopy Platform at the Douglas Hospital Research Centre and we thank Melina Jaramillo Garcia and Dr. Bita Khadivjam for their imaging expertise.

## DATA AVAILABILITY STATEMENT

All data generated for this study are contained within the manuscript. For further queries, the corresponding author NM may be contacted.

## References

1. Allen Institute. (2004). Allen Mouse Brain Atlas. Retrieved April 1, 2020, from Allen Institute for Brain Science website: http://mouse.brainmap.org

2. Alvarez, J. I., Katayama, T., & Prat, A. (2013). Glial influence on the blood brain barrier. Glia, 61(12), 1939–1958. 10.1002/glia.22575

3. Bankhead, P., Loughrey, M. B., Fernández, J. A., Dombrowski, Y., McArt, D. G., Dunne, P. D., McQuaid, S., Gray, R. T., Murray, L. J., Coleman, H. G., James, J. A., Salto-Tellez, M., & Hamilton, P. W. (2017). QuPath: Open source software for digital pathology image analysis. Scientific reports, 7(1), 16878. 10.1038/s41598-017-17204-5

4. Barton, R. A., & Venditti, C. (2014). Rapid evolution of the cerebellum in humans and other great apes. Current biology: CB, 24(20), 2440–2444. 10.1016/j.cub.2014.08.056

5. Batiuk, M. Y., Martirosyan, A., Wahis, J., de Vin, F., Marneffe, C., Kusserow, C., Koeppen, J., Viana, J. F., Oliveira, J. F., Voet, T., Ponting, C. P., Belgard, T. G., & Holt, M. G. (2020). Identification of region-specific astrocyte subtypes at single cell resolution. Nature communications, 11(1), 1220. 10.1038/s41467-019-14198-8

6. Bayraktar, O. A., Bartels, T., Holmqvist, S., Kleshchevnikov, V., Martirosyan, A., Polioudakis, D., Ben Haim, L., Young, A. M. H., Batiuk, M. Y., Prakash, K., Brown, A., Roberts, K., Paredes, M. F., Kawaguchi, R., Stockley, J. H., Sabeur, K., Chang, S. M., Huang, E., Hutchinson, P., Ullian, E. M., … Rowitch, D. H. (2020). Astrocyte layers in the mammalian cerebral cortex revealed by a single-cell in situ transcriptomic map. Nature neuroscience, 23(4), 500–509. 10.1038/s41593-020-0602-1

7. Buckner R. L. (2013). The cerebellum and cognitive function: 25 years of insight from anatomy and neuroimaging. Neuron, 80(3), 807–815. 10.1016/j.neuron.2013.10.044

8. Buckner, R. L., Krienen, F. M., Castellanos, A., Diaz, J. C., & Yeo, B. T. (2011). The organization of the human cerebellum estimated by intrinsic functional connectivity. Journal of neurophysiology, 106(5), 2322–2345. 10.1152/jn.00339.2011

9. Buffo, A., & Rossi, F. (2013). Origin, lineage and function of cerebellar glia. Progress in neurobiology, 109, 42–63. 10.1016/j.pneurobio.2013.08.001

10. Cerrato V. (2020). Cerebellar Astrocytes: Much More Than Passive Bystanders In Ataxia Pathophysiology. Journal of clinical medicine, 9(3), 757. 10.3390/jcm9030757

11. Cerrato, V., Parmigiani, E., Figueres-Oñate, M., Betizeau, M., Aprato, J., Nanavaty, I., Berchialla, P., Luzzati, F., de’Sperati, C., López-Mascaraque, L., & Buffo, A. (2018). Multiple origins and modularity in the spatiotemporal emergence of cerebellar astrocyte heterogeneity. PLoS biology, 16(9), e2005513. 10.1371/journal.pbio.2005513

12. Chambers, T., Escott-Price, V., Legge, S., Baker, E., Singh, K. D., Walters, J. T. R., Caseras, X., & Anney, R. J. L. (2022). Genetic common variants associated with cerebellar volume and their overlap with mental disorders: a study on 33,265 individuals from the UK-Biobank. Molecular psychiatry, 27(4), 2282–2290. 10.1038/s41380-022-01443-8

13. Chung, W.-S., Allen, N.J., Eroglu, C. (2015). Astrocytes control synapse formation, function, and elimination. Cold Spring Harb. Perspect. Biol. 7, a020370. 10.1101/cshperspect.a020370

14. Colombo, J. A., & Reisin, H. D. (2004). Interlaminar astroglia of the cerebral cortex: a marker of the primate brain. Brain research, 1006(1), 126–131. 10.1016/j.brainres.2004.02.003

15. Depping, M. S., Schmitgen, M. M., Kubera, K. M., & Wolf, R. C. (2018). Cerebellar Contributions to Major Depression. Frontiers in psychiatry, 9, 634. 10.3389/fpsyt.2018.00634

16. De Zeeuw, C. I., & Hoogland, T. M. (2015). Reappraisal of Bergmann glial cells as modulators of cerebellar circuit function. Frontiers in cellular neuroscience, 9, 246. 10.3389/fncel.2015.00246

17. Díaz-Castro, B., Robel, S., & Mishra, A. (2023). Astrocyte Endfeet in Brain Function and Pathology: Open Questions. Annual review of neuroscience, 46, 101–121. 10.1146/annurev-neuro-091922-031205

18. Endo, F., Kasai, A., Soto, J. S., Yu, X., Qu, Z., Hashimoto, H., Gradinaru, V., Kawaguchi, R., & Khakh, B. S. (2022). Molecular basis of astrocyte diversity and morphology across the CNS in health and disease. *Science (New York*, N.Y.), 378(6619), eadc9020. 10.1126/science.adc9020

19. Falcone, C., McBride, E. L., Hopkins, W. D., Hof, P. R., Manger, P. R., Sherwood, C. C., Noctor, S. C., & Martínez-Cerdeño, V. (2022). Redefining varicose projection astrocytes in primates. Glia, 70(1), 145–154. 10.1002/glia.24093

20. Falcone, C., Wolf-Ochoa, M., Amina, S., Hong, T., Vakilzadeh, G., Hopkins, W. D., Hof, P. R., Sherwood, C. C., Manger, P. R., Noctor, S. C., & Martínez-Cerdeño, V. (2019). Cortical interlaminar astrocytes across the therian mammal radiation. The Journal of comparative neurology, 527(10), 1654–1674. 10.1002/cne.24605

21. Fastenrath, M., Spalek, K., Coynel, D., Loos, E., Milnik, A., Egli, T., Schicktanz, N., Geissmann, L., Roozendaal, B., Papassotiropoulos, A., & de Quervain, D. J. (2022). Human cerebellum and corticocerebellar connections involved in emotional memory enhancement. Proceedings of the National Academy of Sciences of the United States of America, 119(41), e2204900119. 10.1073/pnas.2204900119

22. Goertzen, A., & Veh, R. W. (2018). Fañanas cells-the forgotten cerebellar glia cell type: Immunocytochemistry reveals two potassium channel-related polypeptides, Kv2.2 and Calsenilin (KChIP3) as potential marker proteins. Glia, 66(10), 2200–2208. 10.1002/glia.23478

23. Haldipur, P., Millen, K. J., & Aldinger, K. A. (2022). Human Cerebellar Development and Transcriptomics: Implications for Neurodevelopmental Disorders. Annual review of neuroscience, 45, 515–531. 10.1146/annurev-neuro-111020-091953

24. Hoogland, T. M., & Kuhn, B. (2010). Recent developments in the understanding of astrocyte function in the cerebellum in vivo. *Cerebellum (London*, England), 9(3), 264–271. 10.1007/s12311-009-0139-z

25. Hwang, I., Kim, B. S., Ko, H. R., Cho, S., Lee, H. Y., Cho, S. W., Ryu, D., Shim, S., & Ahn, J. Y. (2022). Cerebellar dysfunction and schizophrenia-like behavior in Ebp1-deficient mice. Molecular psychiatry, 27(4), 2030–2041. 10.1038/s41380-022-01458-1

26. Karpf, J., Unichenko, P., Chalmers, N., Beyer, F., Wittmann, M. T., Schneider, J., Fidan, E., Reis, A., Beckervordersandforth, J., Brandner, S., Liebner, S., Falk, S., Sagner, A., Henneberger, C., & Beckervordersandforth, R. (2022). Dentate gyrus astrocytes exhibit layer-specific molecular, morphological and physiological features. Nature neuroscience, 25(12), 1626–1638. 10.1038/s41593-022-01192-5

27. Klein, A. P., Ulmer, J. L., Quinet, S. A., Mathews, V., & Mark, L. P. (2016). Nonmotor Functions of the Cerebellum: An Introduction. AJNR. American journal of neuroradiology, 37(6), 1005–1009. 10.3174/ajnr.A4720

28. Korbo, L., & Andersen, B. B. (1995). The distributions of Purkinje cell perikaryon and nuclear volume in human and rat cerebellum with the nucleator method. Neuroscience, 69(1), 151–158. 10.1016/0306-4522(95)00223-6

29. Kozareva, V., Martin, C., Osorno, T., Rudolph, S., Guo, C., Vanderburg, C., Nadaf, N., Regev, A., Regehr, W. G., & Macosko, E. (2021). A transcriptomic atlas of mouse cerebellar cortex comprehensively defines cell types. Nature, 598(7879), 214–219. 10.1038/s41586-021-03220-z

30. Kreutz, A., & Barger, N. (2018). Maximizing Explanatory Power in Stereological Data Collection: A Protocol for Reliably Integrating Optical Fractionator and Multiple Immunofluorescence Techniques. Frontiers in neuroanatomy, 12, 73. 10.3389/fnana.2018.00073

31. Krupenko S. A. (2009). FDH: an aldehyde dehydrogenase fusion enzyme in folate metabolism. Chemico-biological interactions, 178(1-3), 84–93. 10.1016/j.cbi.2008.09.007

32. Lange W. (1975). Cell number and cell density in the cerebellar cortex of man and some other mammals. Cell and tissue research, 157(1), 115–124. 10.1007/BF00223234

33. Madigan, J. C., & Carpenter, M. B. (1971). Cerebellum of the rhesus monkey: atlas of lobules, laminae, and folia, in sections. University Park Press.

34. Madisen, L., Zwingman, T. A., Sunkin, S. M., Oh, S. W., Zariwala, H. A., Gu, H., Ng, L. L., Palmiter, R. D., Hawrylycz, M. J., Jones, A. R., Lein, E. S., & Zeng, H. (2010). A robust and high-throughput Cre reporting and characterization system for the whole mouse brain. Nature neuroscience, 13(1), 133–140. 10.1038/nn.2467

35. Masoli, S., Sanchez-Ponce, D., Vrieler, N. et al. Human Purkinje cells outperform mouse Purkinje cells in dendritic complexity and computational capacity. Commun Biol 7, 5 (2024). 10.1038/s42003-023-05689-y

36. Matias, I., Morgado, J., & Gomes, F. C. A. (2019). Astrocyte Heterogeneity: Impact to Brain Aging and Disease. Frontiers in aging neuroscience, 11, 59. 10.3389/fnagi.2019.00059

37. Mayorquin, L. C., Rodriguez, A. V., Sutachan, J. J., & Albarracín, S. L. (2018). Connexin-Mediated Functional and Metabolic Coupling Between Astrocytes and Neurons. Frontiers in molecular neuroscience, 11, 118. 10.3389/fnmol.2018.00118

38. Miyazaki, T., Yamasaki, M., Hashimoto, K., Kohda, K., Yuzaki, M., Shimamoto, K., Tanaka, K., Kano, M., & Watanabe, M. (2017). Glutamate transporter GLAST controls synaptic wrapping by Bergmann glia and ensures proper wiring of Purkinje cells. Proceedings of the National Academy of Sciences of the United States of America, 114(28), 7438–7443. 10.1073/pnas.1617330114

39. Mouton, P.R. (2002). Principles and Practices of Unbiased Stereology. Baltimore: The Johns Hopkins University Press.

40. Muñoz, Y., Cuevas-Pacheco, F., Quesseveur, G., & Murai, K. K. (2021). Light microscopic and heterogeneity analysis of astrocytes in the common marmoset brain. Journal of neuroscience research, 99(12), 3121–3147. 10.1002/jnr.24967

41. Oberheim, N. A., Wang, X., Goldman, S., & Nedergaard, M. (2006). Astrocytic complexity distinguishes the human brain. Trends in neurosciences, 29(10), 547–553. 10.1016/j.tins.2006.08.004

42. Oberheim, N. A., Takano, T., Han, X., He, W., Lin, J. H., Wang, F., Xu, Q., Wyatt, J. D., Pilcher, W., Ojemann, J. G., Ransom, B. R., Goldman, S. A., & Nedergaard, M. (2009). Uniquely hominid features of adult human astrocytes. The Journal of neuroscience : the official journal of the Society for Neuroscience, 29(10), 3276–3287. 10.1523/JNEUROSCI.4707-08.2009

43. O’Leary, L. A., Davoli, M. A., Belliveau, C., Tanti, A., Ma, J. C., Farmer, W. T., Turecki, G., Murai, K. K., & Mechawar, N. (2020). Characterization of Vimentin-Immunoreactive Astrocytes in the Human Brain. Frontiers in neuroanatomy, 14, 31. 10.3389/fnana.2020.00031

44. Romer, A. L., Knodt, A. R., Houts, R., Brigidi, B. D., Moffitt, T. E., Caspi, A., & Hariri, A. R. (2018). Structural alterations within cerebellar circuitry are associated with general liability for common mental disorders. Molecular psychiatry, 23(4), 1084–1090. 10.1038/mp.2017.57

45. Schmahmann J. D. (2019). The cerebellum and cognition. Neuroscience letters, 688, 62–75. 10.1016/j.neulet.2018.07.005

46. Schmahmann, J.D., Doyon, J., Toga, A.W., Petrides, M., & Evans, A.C. (2000). MRI Atlas of the Human Cerebellum. Academic Press.

47. Sereno, M. I., Diedrichsen, J., Tachrount, M., Testa-Silva, G., d’Arceuil, H., & De Zeeuw, C. (2020). The human cerebellum has almost 80% of the surface area of the neocortex. Proceedings of the National Academy of Sciences of the United States of America, 117(32), 19538–19543. 10.1073/pnas.2002896117

48. Shibuki, K., Gomi, H., Chen, L., Bao, S., Kim, J. J., Wakatsuki, H., Fujisaki, T., Fujimoto, K., Katoh, A., Ikeda, T., Chen, C., Thompson, R. F., & Itohara, S. (1996). Deficient cerebellar long-term depression, impaired eyeblink conditioning, and normal motor coordination in GFAP mutant mice. Neuron, 16(3), 587–599. 10.1016/s0896-6273(00)80078-1

49. Srinivasan, R., Lu, T. Y., Chai, H., Xu, J., Huang, B. S., Golshani, P., Coppola, G., & Khakh, B. S. (2016). New Transgenic Mouse Lines for Selectively Targeting Astrocytes and Studying Calcium Signals in Astrocyte Processes In Situ and In Vivo. Neuron, 92(6), 1181–1195. 10.1016/j.neuron.2016.11.030

50. Stogsdill, J. A., Harwell, C. C., & Goldman, S. A. (2023). Astrocytes as master modulators of neural networks: Synaptic functions and disease-associated dysfunction of astrocytes. Annals of the New York Academy of Sciences, 1525(1), 41–60. 10.1111/nyas.15004

51. Sosunov, A. A., Wu, X., Tsankova, N. M., Guilfoyle, E., McKhann, G. M., 2nd, & Goldman, J. E. (2014). Phenotypic heterogeneity and plasticity of isocortical and hippocampal astrocytes in the human brain. The Journal of neuroscience : the official journal of the Society for Neuroscience, 34(6), 2285–2298. 10.1523/JNEUROSCI.4037-13.2014

52. Verkhratsky, A., Parpura, V., Li, B., & Scuderi, C. (2021). Astrocytes: The Housekeepers and Guardians of the CNS. Advances in neurobiology, 26, 21–53. 10.1007/978-3-030-77375-5_2

53. Viana, J. F., Machado, J. L., Abreu, D. S., Veiga, A., Barsanti, S., Tavares, G., Martins, M., Sardinha, V. M., Guerra-Gomes, S., Domingos, C., Pauletti, A., Wahis, J., Liu, C., Calì, C., Henneberger, C., Holt, M. G., & Oliveira, J. F. (2023). Astrocyte structural heterogeneity in the mouse hippocampus. Glia, 71(7), 1667–1682. 10.1002/glia.24362

54. Wilhelmsson, U., Li, L., Pekna, M., Berthold, C. H., Blom, S., Eliasson, C., Renner, O., Bushong, E., Ellisman, M., Morgan, T. E., & Pekny, M. (2004). Absence of glial fibrillary acidic protein and vimentin prevents hypertrophy of astrocytic processes and improves post-traumatic regeneration. The Journal of neuroscience : the official journal of the Society for Neuroscience, 24(21), 5016–5021. 10.1523/JNEUROSCI.0820-04.2004

55. Yamada, K., Fukaya, M., Shibata, T., Kurihara, H., Tanaka, K., Inoue, Y., & Watanabe, M. (2000). Dynamic transformation of Bergmann glial fibers proceeds in correlation with dendritic outgrowth and synapse formation of cerebellar Purkinje cells. The Journal of comparative neurology, 418(1), 106–120.

56. Yang, Y., Vidensky, S., Jin, L., Jie, C., Lorenzini, I., Frankl, M., & Rothstein, J. D. (2011). Molecular comparison of GLT1+ and ALDH1L1+ astrocytes in vivo in astroglial reporter mice. Glia, 59(2), 200–207. 10.1002/glia.21089

57. Yang, Z., & Wang, K. K. (2015). Glial fibrillary acidic protein: from intermediate filament assembly and gliosis to neurobiomarker. Trends in neurosciences, 38(6), 364–374. 10.1016/j.tins.2015.04.003

